# Visual Attention in Peripersonal Space

**DOI:** 10.1101/2025.10.20.683534

**Authors:** Hamidreza Ramezanpour, Devin Heinze Kehoe, Carolyn J. Perry, Mazyar Fallah

## Abstract

Bringing the hand near a visual stimulus enhances visual processing. This effect is linked to peripersonal space (PPS), the body-centered region where visual and proprioceptive information interact. Despite extensive behavioral evidence, the neural basis of this interaction in early visual cortex remains unclear. In this study, we investigated how hand proximity modulates orientation selectivity in area V2 by recording single-neuron responses from two rhesus monkeys. The monkeys performed a fixation task while their hand was positioned near a visual stimulus while either being visible or occluded, and compared with when the hand was away from the stimulus. When the near hand was visible, neural firing rates in V2 were significantly higher, accompanied by sharper orientation tuning. In contrast, occluding the hand broadened orientation tuning compared to when the hand was away. These effects emerged rapidly after stimulus onset and were coherent across the population, demonstrating that PPS is actively prioritized during visual processing. Together, the findings reveal two complementary (feedback) signals in V2: a congruence-driven enhancement when visual and proprioceptive inputs align, and a mismatch-driven suppression when they conflict, indicating that V2 integrates multisensory cues to encode PPS and support action-relevant visual processing.

## Introduction

The phenomenon of enhanced visual processing near the hand has attracted considerable attention in recent years, with numerous studies documenting improvements in various visual tasks when a hand is placed close to the stimulus. For example, it has been shown that target detection and figure-ground discrimination are faster and more accurate when the hand is near the visual stimulus (Reed et al., 2006, 2010; Jackson et al., 2010). These enhancements extend to more complex processes like working memory, where stimuli located near the hand are remembered more effectively (Tseng and Bridgeman, 2011). Other studies have demonstrated that this near-hand effect improves orientation processing, which is critical for tasks involving reaching and grasping (Craighero et al., 1999; Hannus et al., 2005; Brown et al., 2008). These behavioral results suggest that the hand’s proximity modulates visual processing in a way that supports more efficient motor actions, potentially driven by specialized neural mechanisms in the brain.

One proposed explanation for this effect is that near-hand space is subject to enhanced attentional selection (di Pellegrino G and Frassinetti, 2000; Schendel and Robertson, 2004; Reed et al., 2006, 2010; Abrams et al., 2008; Brown et al., 2008). According to this view, one can hypothesize that fronto-parietal bimodal neurons that process both visual and somatosensory inputs play a role in boosting attentional resources allocated to the space around the hand via feedback mechanisms (Perry et al., 2016; Perry and Fallah, 2017). This view gets support from the fact that there are direct projections from parietal areas such as AIP and V6A that are involved in tactile object recognition (Rizzolatti and Matelli, 2003) and grasping (Fattori et al., 2009, 2010) to visual areas such as V2 (Passarelli et al., 2011). V2 is known to be modulated by attention and is critical for processing features such as orientation and shape (Motter, 1993; Luck et al., 1997). Perry and colleagues provided direct neurophysiological evidence showing that the presence of a nearby hand enhances the orientation selectivity of neurons in area V2, leading to sharper tuning curves and reduced response variability (Perry et al., 2015). This sharpening of orientation tuning could potentially facilitate more precise discrimination of action-relevant features, which would aid in reaching and grasping tasks.

Despite these advances, the sources of the enhanced visual response near the hand remain unclear. The current experiment aims to test three distinct hypotheses: H1, that proprioception alone enhances visual processing near the hand; H2, that visual attention is the primary driver of the effect; and H3, that both proprioceptive and visual signals contribute to the enhancement. By distinguishing between these hypotheses, we hope to clarify the sensory processes that drive the enhanced visual processing near the hand and provide a more comprehensive understanding of how the brain integrates sensory inputs to optimize actions in peripersonal space (PPS).

## Materials and Methods

Two adult male rhesus monkeys were used in this study. The monkeys were surgically implanted with a head post and a recording chamber over the left visual area V2, using stereotaxic coordinates. The placement of the recording chamber was verified by receptive field mapping and the topographic organization of V2, as described in our previous work (Perry et al., 2015). A tungsten electrode was advanced using a microdrive (Crist Instruments), and neuronal data were recorded with a Multichannel Acquisition Processor (Plexon Inc.). Neurons were isolated using Rasputin software during recording, and the receptive fields were mapped using a manually controlled flashing bar that varied in size, orientation, and position. Neurons were isolated offline using Offline Sorter (Plexon Inc.) for subsequent analyses. All experimental and surgical procedures complied with animal care guidelines as defined by the CACC (Canadian Animal Care Committee) and York University’s Animal Care Committee.

### Experimental Design

The monkeys were trained to fixate on a point displayed on a monitor placed 36 cm away while reaching with their right hand to grasp a vertically oriented touch bar next to the monitor (**Figure 1**). To prevent interference with baseline neural activity, the distance between the touch bar and the visual receptive field was carefully controlled, ensuring that the monkey’s hand remained outside the receptive field. Three conditions were tested: in the Hand-Near condition, the monkey grasped the touch bar, positioning its hand near the visual stimulus but still outside the receptive field; in the Hand-Away condition, the monkey did not reach for the touch bar, keeping its hand away from the stimulus; and in the Hand-Occluded condition, the hand was placed near the stimulus but covered with a black shield, preventing the monkey from seeing it while still maintaining the grasp. During each trial, a fixation point appeared, and once fixation was maintained, an oriented rectangle was presented within the visual receptive field for 400 ms. The rectangle’s orientation varied (0º to 157.5º), and each orientation was repeated 10-20 times in each condition. The monkey was rewarded for maintaining both fixation and contact with the touch bar throughout the trial. This paradigm was designed to replicate our prior study (Perry et al., 2015) while adding the Hand-Occluded condition to separate the visual and proprioceptive contributions to changes in neuronal responses in V2.

**Figure 1.**
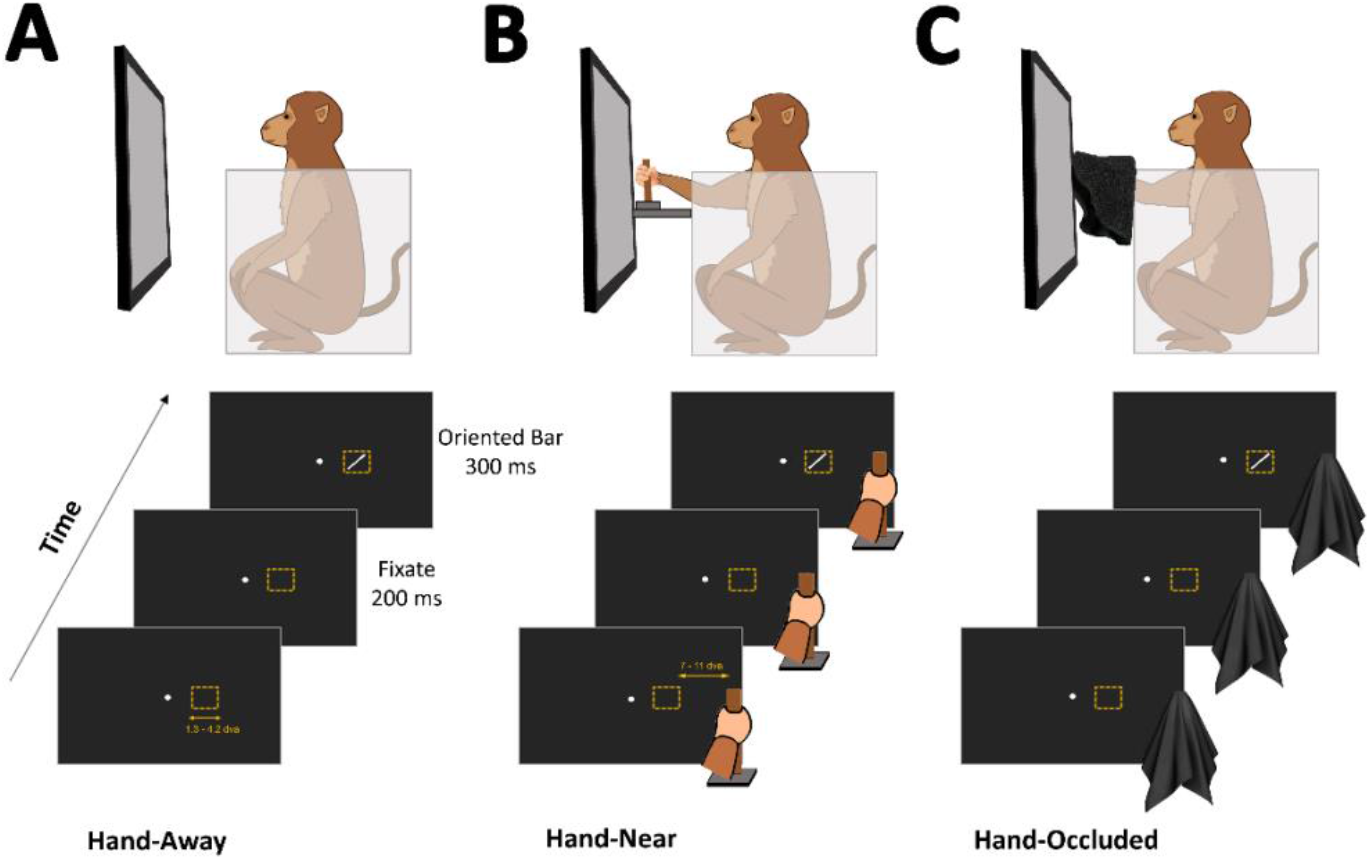
Experimental setup. Three conditions were tested: (A) Hand-Away – the monkey’s hand was kept away from the touch bar. (B) Hand-Near – the monkey grasped the touch bar, positioning its hand close to the visual stimulus but still outside the receptive field. (C) Hand-Occluded – the monkey’s hand was positioned near the stimulus but shielded from view, allowing for grasping without visual feedback. In each trial, the monkey fixated on a small fixation point for 200 ms before a brief (400 ms) display of an oriented bar within the receptive field. Different orientations were shown repeatedly across conditions, and the monkey was rewarded for maintaining both fixation and contact with the touch bar.

### Data Analysis

Neuronal responses were analyzed from -175 ms prior to stimulus onset (baseline) to 400 ms post-stimulus. A one-way ANOVA was used to select neurons that had significantly higher firing rates for at least one of the 8 oriented for further analysis (n=67 neurons). We calculated the orientation tuning modulation index for each neuron as:

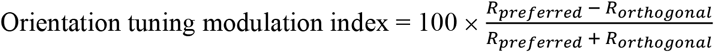

in which *R*_*preferred*_ and *R*_*orthogonal*_ are the neuron’s response rates to preferred orientation (largest response) and the orientation that is 90 degrees away from it. Statistical significance between different conditions was evaluated using the Wilcoxon signed-rank test.

Principal Component Analysis (PCA) was applied to neural response data across three conditions (Hand-Near, No-Hand, and Hand-Occluded) to reduce dimensionality and extract the main patterns of temporal variance. We analyzed direction-averaged, baseline-subtracted firing rates (baseline: −175–0 ms; 0 ms = stimulus onset) for the three conditions. To compare neurons on a common scale and down-weight trivial global gain, we z-scored each neuron across all time points and conditions (unit variance), then applied PCA to the concatenated time×neuron matrix. The first three principal components, representing the dominant trends in neuronal activity, were analyzed over time to compare response dynamics between conditions. To assess when conditions diverge in low-dimensional space, we performed paired Hotelling’s T^2^ on PC1–PC3 using non-overlapping 20 ms bins.

We also quantified condition information (Hand-Near, Hand-Away, Hand-Occluded) with a 3-class leave-one-direction-out (LODO) decoder on neural activities. For each neuron and direction, we binned the time series into non-overlapping 20 ms bins. Within each bin we averaged firing rates, yielding one population vector per direction and condition. For each time bin, we performed LODO over the 8 directions: train on the 7 remaining directions and test on the held-out direction (3 test items: Hand-Near, Hand-Away, Hand-Occluded). Features were z-scored using training statistics (per fold); zero-variance features were removed. The classifier was LDA (diagLinear) with small regularization (Gamma = 0.01), with a nearest-centroid (cosine) fallback if LDA failed to train. Bin accuracy was the mean across the 8 folds. For each bin we also computed a 95% bootstrap CI by resampling the LODO folds (directions) with replacement (B = 1000 bootstrap replicates) and taking the 2.5th/97.5th percentiles of the mean accuracy. To test for statistical significance testing, we ran a permutation test with R = 1000 label shuffles applied only to the training labels (test labels left intact) within each LODO fold, recomputing accuracy each time to form a null distribution. The one-sided p-value was the fraction of null accuracies ≥ the observed accuracy. To control for testing across bins, we applied Benjamini–Hochberg FDR at q = 0.05 across time and reported significant time spans as contiguous bins passing the FDR threshold.

## Results

We observed a diverse pattern of neural responses across neurons and conditions. **Figure 2A**, shows a pure visual neuron that did not show any significant modulations when the hand was close to the visual stimulus. **Figure 2B**, shows a neuron for which the response to the preferred orientation was modulated by the presence of the hand with no significant difference when hand was visible or occluded. Figure **2C**, shows a neuron that was not initially tuned in the Hand-Away condition, but became tuned when a visible hand was placed near the stimulus. **Figure 2D**, shows a neuron for which the presence of the hand leads to an increase in firing to the nonpreferred orientations. For the example neuron in **Figure 2E**, occlusion of the hand reduced the firing, especially to the preferred orientation. And, for the neuron in **Figure 2F**, a visible hand near the stimulus enhanced the firing, especially to the preferred orientation.

**Figure 2.**
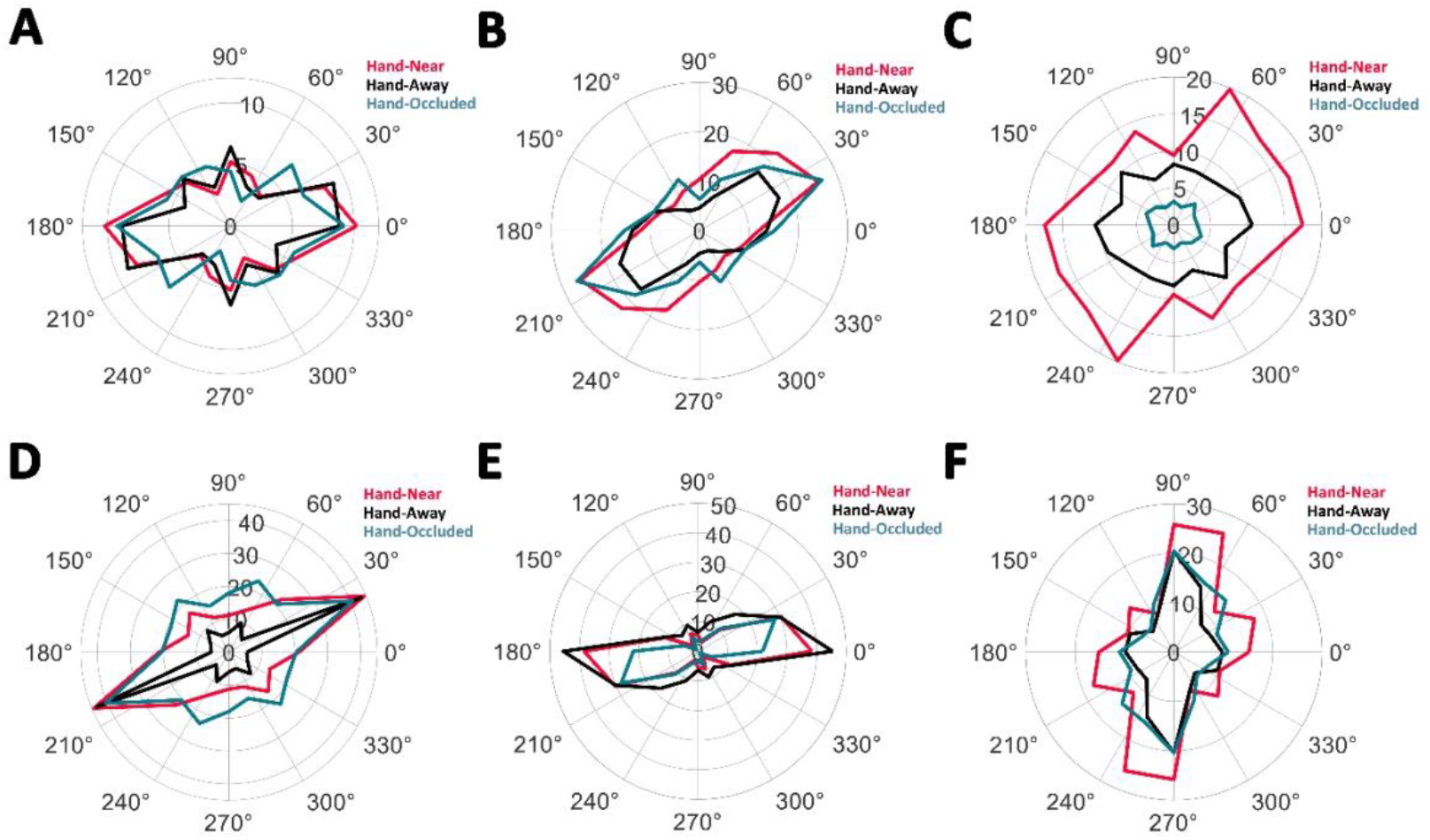
Neural response diversity across conditions. Polar plots illustrate examples of neurons with distinct response patterns based on hand proximity and visibility. (A) A neuron exhibiting purely visual responses with no modulation by hand proximity. (B) A neuron showing modulation of responses to the preferred orientation by hand presence, regardless of hand visibility. (C) A neuron acquiring stimulus tuning in the Hand-Near condition with a visible hand. (D) A neuron with increased firing to nonpreferred orientations when the hand is present. (E) A neuron demonstrating reduced firing, particularly to the preferred orientation, when the hand is occluded. (F) A neuron showing enhanced firing to the preferred orientation when a visible hand is near the stimulus.

**Figure 3A** shows the average population response across the three conditions—Hand-Near (red), Hand-Away (black), and Hand-Occluded (blue)—aligned to stimulus onset (time = 0 ms). Hand-Near consistently evokes the highest firing rates, peaking shortly after onset (~150 ms) and remaining elevated throughout the analysis window. Hand-Away produces a significantly lower response (Wilcoxon signed-rank, p < 0.001), with a smaller peak. Hand-Occluded is intermediate: firing rates are higher than Hand-Away (p < 0.01, Wilcoxon signed-rank) and not significantly lower than Hand-Near (p > 0.05; see right panel of **Figure. 3A**). **Figure 3B** quantifies orientation selectivity via a modulation index (larger values indicate sharper tuning). Average tuning modulation is highest in the Hand-Near condition (p < 0.05), is significantly reduced in Hand-Away (p < 0.05, Wilcoxon signed-rank), and is lowest in Hand-Occluded (p < 0.05, Wilcoxon signed-rank), indicating that proximity of the hand increases orientation selectivity whereas occlusion broadens tuning. Condition decoding from population activity (leave-one-direction-out across directions) rises above chance (~33%) by ~50 ms after stimulus onset (**Figure. 3C**), showing that hand context is linearly readable remarkably early. Together with the divergences in PSTHs and tuning, this indicates that those effects converge into a robust condition signal that a simple linear downstream readout could exploit to represent PPS.

**Figure 3.**
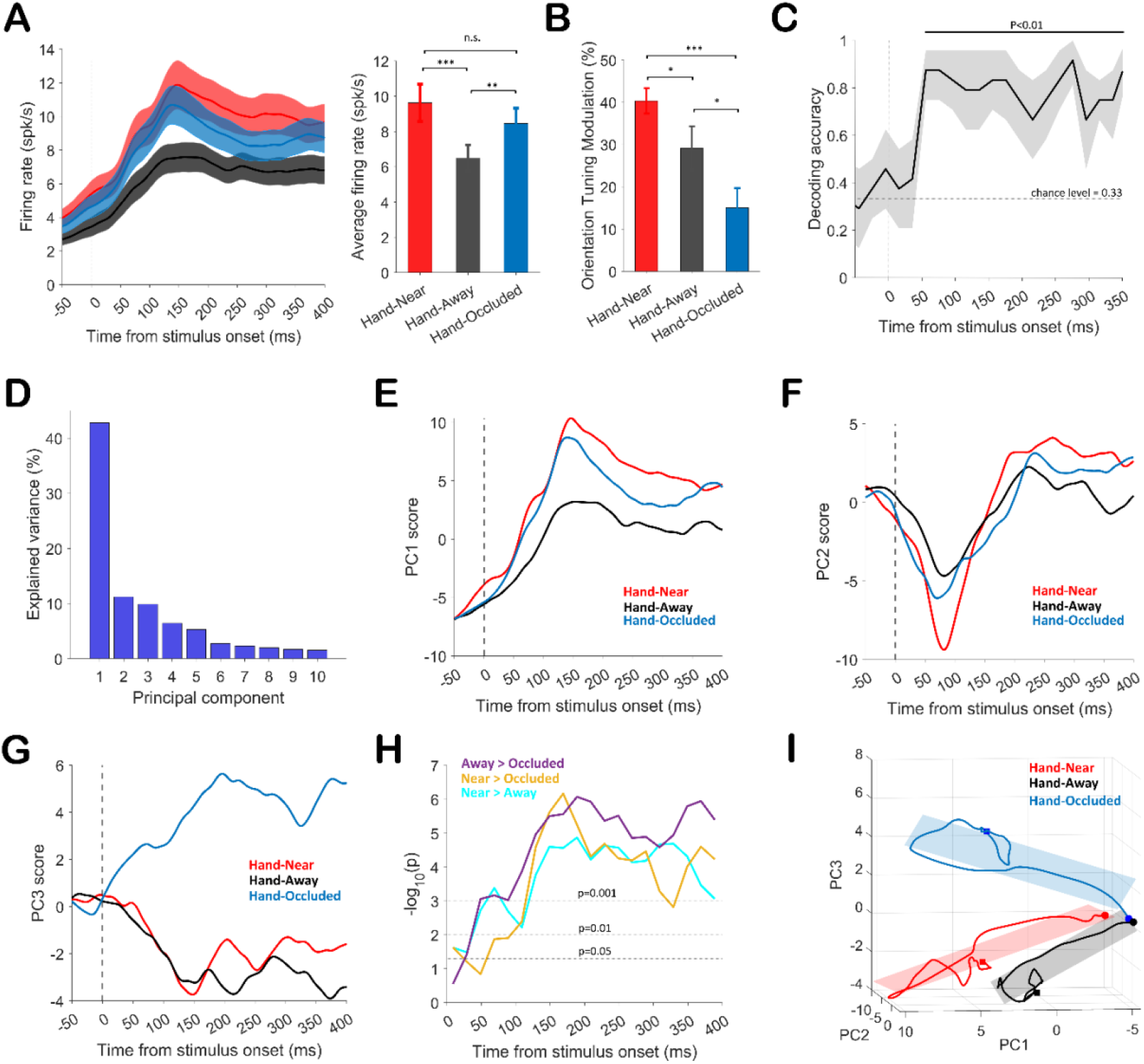
Neural population responses and principal component analysis (PCA) reveal distinct effects of hand proximity and visibility. Color code for all panels: Hand-Near (red), Hand-Away (black), Hand-Occluded (blue). (A) Population peri-stimulus time histograms (PSTHs) aligned to stimulus onset (0 ms). Hand-Near shows the largest response, Hand-Away the smallest, and Hand-Occluded intermediate. (B) Orientation-tuning modulation index: tuning is sharpest in Hand-Near, reduced in Hand-Away, and lowest in Hand-Occluded. (C) Condition decoding (leave-one-direction-out) rises above chance (~33%) by ~50 ms after onset, indicating early linear readout of hand context. (D) PCA variance spectrum; PC1 explains 42.78% of variance. (E–G) Time courses of PC1–PC3 computed from activity in a 400-ms post-onset window: PC1 tracks overall response magnitude (peak ~150 ms; Hand-Near > Hand-Occluded > Hand-Away), PC2 captures earlier temporal dynamics (troughs ~70–80 ms earlier than PC1), and PC3 highlights divergence under occlusion (late positive phase ~200 ms). (H) Time bins (20 ms, non-overlapping) with significant between-condition differences in PC trajectories (paired Hotelling’s T^2^). (I) Low-dimensional trajectory geometry (PC1–PC3): Hand-Near and Hand-Away lie in near-coplanar subspaces, whereas Hand-Occluded diverges; circle marks stimulus onset and square marks stimulus duration endpoint (onset + 400 ms). Error bars indicate SEM. Statistical notation: p < 0.05 (), p < 0.01 (), p < 0.001 (), and n.s. indicates not significant.

We next examined the geometry of the population response with PCA across Hand-Near, Hand-Away, and Hand-Occluded. The explained-variance spectrum was strongly front-loaded, with PC1 alone capturing 42.78% of the variance (**Figure. 3D**), implying that condition information is concentrated along a small number of axes and is therefore amenable to efficient linear readout. **Figures 3E–G** show PC time courses derived from projections of the average responses within a 400-ms post-stimulus window. PC1 primarily tracks overall response magnitude: all three conditions share a clear peak around ~150 ms, with Hand-Near > Hand-Occluded > Hand-Away, paralleling the PSTHs in **Figure 3A**. PC2 captures similar dynamics but shifted earlier in time, with troughs occurring ~70–80 ms before those peaks in PC1 (**Figure. 3F**). PC3 highlights variance associated with occlusion: the Hand-Occluded condition shows a more prominent late positive phase peaking around ~200 ms compared to the other conditions (**Figure. 3G**). Time windows with significant differences between PC trajectories were identified using a paired Hotelling’s T^2^ test on non-overlapping 20-ms bins (**Figure. 3H**). Higher PCs explained progressively less variance (**Figure. S1**). Finally, principal-component trajectories (PC1–PC3) revealed distinct population geometries (**Figure. 3I**). Planes fit to post-stimulus activity showed that Hand-Near and Hand-Away were nearly coplanar (10.1° between plane normals), whereas Hand-Occluded diverged into a different subspace (31.2° from Hand-Near; 41.0° from Hand-Away). Trajectory lengths also differed (Hand-Near > Hand-Occluded > Hand-Away), indicating the strongest temporal evolution when the hand was visible, intermediate dynamics under occlusion, and minimal modulation when the hand was absent (**Figure S2**). Together, these results suggest that Hand-Near and Hand-Away share a common low-dimensional manifold, while occlusion engages a more orthogonal representational mode; the magnitude of trajectory length reflects the strength of neural encoding within each subspace. Finally, significant correlations emerged between PCs, especially higher order ones, and orientation tuning modulation indices (OTM), indicating that specific dimensions of the population activity encode orientation tuning in a condition-dependent manner (**Figure S3**). While some PCs (e.g., PC2, PC4, PC6) showed negative correlations with OTM, others (PC5, PC8) showed positive correlations, suggesting that different subspaces differentially capture orientation information across hand contexts.

## Discussion

In this study, we provide compelling evidence that the proximity of the hand modulates visual processing in the early visual cortex, area V2. Our findings build upon previous research documenting alterations in visual processing near the hand. Numerous behavioral studies have shown that visual processing is enhanced when the hand is close to a visual stimulus (Craighero et al., 1999; Hannus et al., 2005; Reed et al., 2006, 2010; Jackson et al., 2010). This enhancement has been hypothesized to arise from attentional mechanisms that prioritize near-hand space, potentially mediated by fronto-parietal bimodal neurons (di Pellegrino G and Frassinetti, 2000; Abrams et al., 2008).

Our study extends these findings by explicitly dissociating the contributions of visual and proprioceptive signals to neural activity in area V2. We examined responses under three conditions: Hand-Near, where both visual and proprioceptive information about the hand are available; Hand-Occluded, where the hand is positioned near the stimulus but hidden from view, isolating proprioceptive input; and Hand-Away, where neither visual nor proprioceptive signals are present in PPS. We found that Hand-Near replicated prior reports, with significantly higher firing rates and sharper orientation tuning compared to Hand-Away (Perry et al., 2015). Importantly, the Hand-Occluded condition yielded intermediate responses on firing rates, demonstrating that proprioception alone can enhance visual responses, but that the full effect requires the joint contribution of visual and proprioceptive signals. This graded pattern highlights the multisensory nature of the effect and supports the idea that PPS is represented in early visual cortex through the integration of both sensory modalities. Surprisingly, we found that occluding vision of the hand actively impaired orientation selectivity: tuning was significantly broader than in the Hand-Away condition. The mismatch between vision and proprioception of the hand impairing visual processing in PPS suggests that both sensory modalities are necessary to encode PPS in the visual system.

Our decoding results further demonstrate that these effects emerge remarkably early in V2. As soon as ~50 ms after stimulus onset, a simple linear decoder could reliably dissociate the three hand conditions, indicating that the combined influence of vision and proprioception converges into a robust condition signal that is readily available to downstream areas. In this scenario, V2 itself could act as a driver, providing early, linearly accessible information about hand context to higher-order regions involved in PPS processing. At the same time, the block-design nature of our task, in which the hand position remained fixed throughout each block, means that high-level feedback—whether visual or proprioceptive or a combination of both—was present even before stimulus onset. Thus, the ~50 ms latency likely reflects the time required for V2 neurons to respond to the visual stimulus in their receptive fields and to integrate this with the continuously available hand-related signals, producing a condition-specific separation at the population level. Therefore, hand position in PPS is an active signal continuously provided to visual processing.

The distinct temporal dynamics captured by our PCA suggest that multiple sources of feedback shape V2 activity depending on the availability of visual and proprioceptive information about the hand. Prior work has identified three classes of parietal regions that could provide such feedback: those dominated by visual representations of the hand (e.g., area V6A (Breveglieri et al., 2016; Galletti et al., 2022)), those with a predominance of proprioceptive hand-related signals (e.g., area PEc and medial intraparietal cortex (Breveglieri et al., 2006)), and those containing bimodal neurons that integrate both modalities (such as anterior intraparietal area, AIP, and ventral intraparietal area, VIP) (Avillac et al., 2005, 2007; Fattori et al., 2009, 2010).

Our decoding analysis showed that condition information emerges as early as ~50 ms after stimulus onset, implying that high-level hand-related feedback—either visual, proprioceptive, or both—is already available in V2 before or immediately upon stimulus arrival. The different principal components then emphasize distinct temporal signatures of this integration. PC1 largely tracks response gain and peaks around 150 ms, consistent with a strong visual drive modulated by hand context. PC2 captured dynamics similar to PC1 but shifted earlier in time, suggesting that distinct inputs may contribute to hand-related modulation in V2. One possibility is that PC2 reflects an additional channel of input, such as proprioceptive signals from parietal circuits encoding hand position, which are available even before stimulus onset. PC3, by contrast, highlights a late divergence in the Hand-Occluded condition (~200 ms), consistent with bimodal or integrative feedback that combines visual and proprioceptive information but is specifically sensitive to mismatches between them. Together, these dynamics suggest that multiple parietal channels converge onto V2: visual-only regions may reinforce the gain of hand-near responses, proprioceptive-dominated areas may contribute to the earliest divergences in the population signal, and bimodal regions may shape later, condition-specific trajectories. Unlike feedback from the frontal eye fields, which enhances visual responses at the target of impending saccades (Moore and Fallah, 2001, 2004; Moore and Armstrong, 2003), our findings suggest that hand-related modulation in V2 arises from two distinct feedback signals.

The first resembles oculomotor feedback: when visual and proprioceptive information about the hand are congruent, as in the Hand-Near condition, V2 neurons show enhanced firing rates and sharper orientation tuning. This enhancement likely reflects an effector-specific gain mechanism that prioritizes stimuli near the hand, analogous to saccade-related enhancement of visual responses at oculomotor targets.

The second signal emerges when visual and proprioceptive inputs are incongruent, as in the Hand-Occluded condition. Here, reciprocal feedback from parietal regions involved in sensorimotor integration (Passarelli et al., 2011; Markov et al., 2014) likely transmits a mismatch signal that transiently impairs orientation selectivity. This suggests that V2 not only integrates multisensory information about the hand but also detects discrepancies between modalities, dynamically adjusting its tuning when visual feedback does not match proprioceptive expectations.

This mismatch mechanism may also help explain everyday behaviors in which people instinctively minimize sensory conflict. For example, when using a tool or manipulating an object that is out of view, such as tightening a screw behind a device or reaching under a sink, people often close their eyes. By removing unreliable visual input, the brain restores consistency between visual and proprioceptive signals, reducing interference and improving control. Although anecdotal, such behaviors illustrate how the visual system may actively seek congruence between modalities, echoing the neural dynamics observed here in V2.

These two forms of feedback, one enhancing under congruence and one suppressive under mismatch, reveal a more complex organization of hand-related modulation than previously recognized. Prior work (Perry et al., 2016) described effector-based attention as a single mechanism enhancing visual processing near the hand; our results demonstrate that it actually comprises two interacting components: a gain-enhancing congruence signal and a suppressive mismatch signal mediated by reciprocal parietal connections.

The results therefore suggest that near-hand attention arises from a distinct, effector-based mechanism that operates alongside, but separately from, traditional oculomotor-driven spatial attention (Moore and Fallah, 2001, 2004). This mechanism may serve to prioritize visual information relevant for grasping and other motor actions, improving orientation selectivity in V2 when cues are congruent, but attenuating it when sensory feedback is inconsistent. Together, these findings provide evidence that both proprioceptive and visual feedback actively contribute to PPS modulation through recurrent feedback loops linking parietal and visual areas.

In terms of future directions, our results open up several intriguing questions. First, it would be valuable to explore whether similar effects are observed in other early visual areas, such as V1 or V4, to determine the extent of this modulation across the visual hierarchy. Additionally, given behavioral and TMS evidence that hand and pointing movements modulate visual orientation processing (Neggers and Bekkering, 2001, 2002; Gutteling et al., 2013) (e.g., Neggers et al., 2006; Sober & Sabes, 2003), investigating how these mechanisms operate during dynamic hand movements—as opposed to static hand positions—would provide further insight into how sensory feedback is integrated during real-time actions. Finally, understanding how these mechanisms are altered in conditions such as motor control disorders could lead to targeted interventions that enhance sensorimotor integration and improve functional outcomes for affected individuals.

## Supplementary Figures

**Figure S1.**
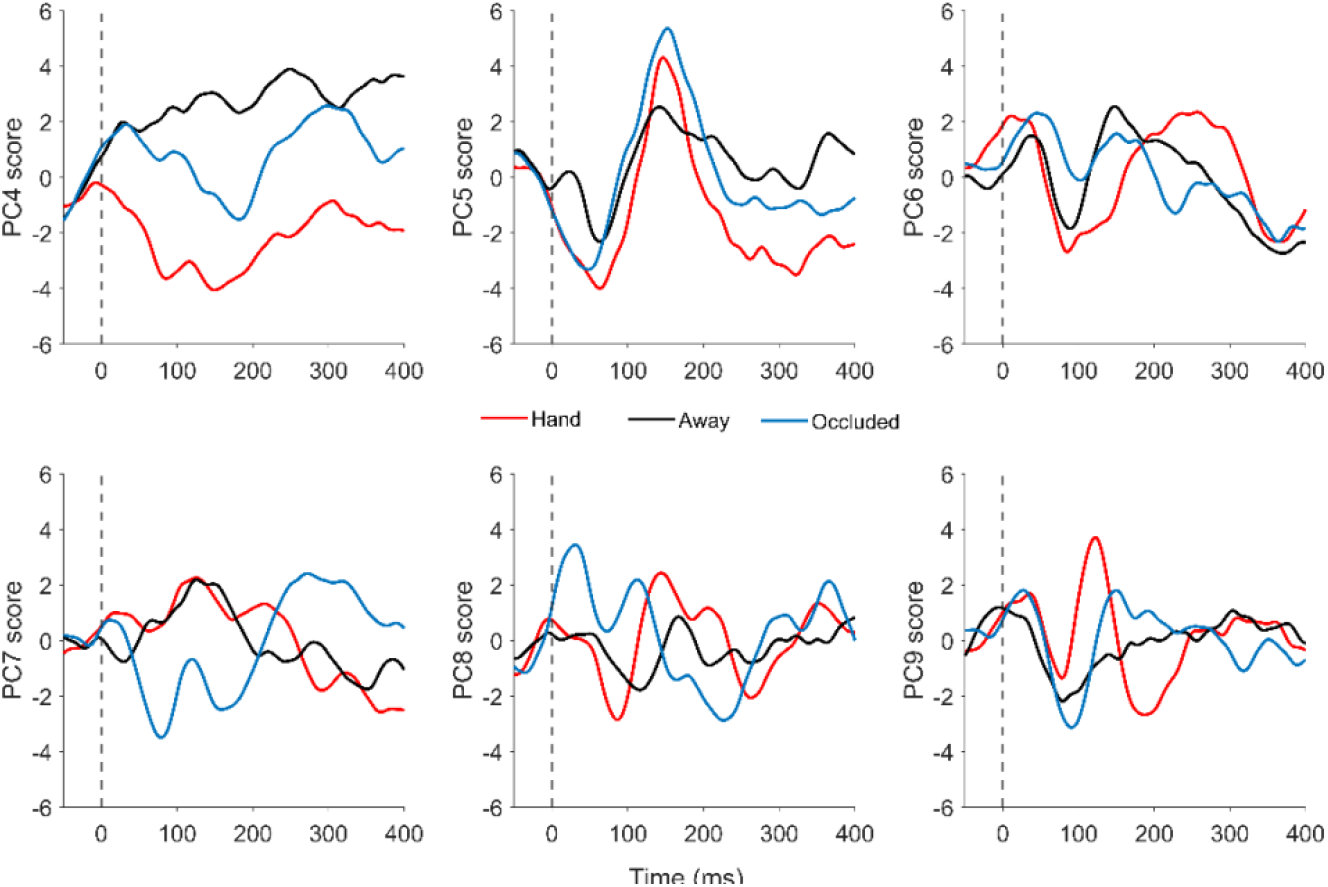
Temporal time courses of principal components PC4 through PC10 computed from population activity in the 0–400 ms post-stimulus window, aligned to stimulus onset (0 ms).

**Figure S2.**
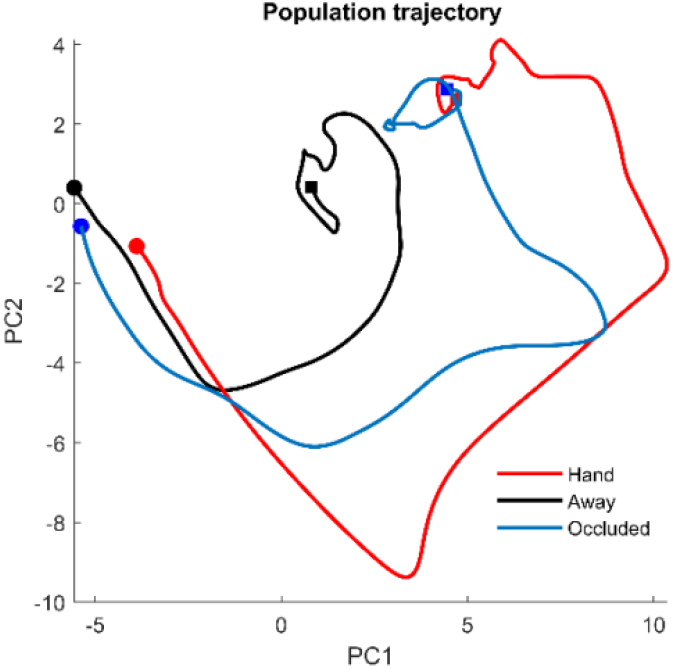
Low-dimensional population trajectory geometry across hand conditions. Visible hand proximity evokes the strongest temporal evolution of neural states, occlusion drives intermediate dynamics, and absence of the hand produces minimal modulation.

**Figure S3.**
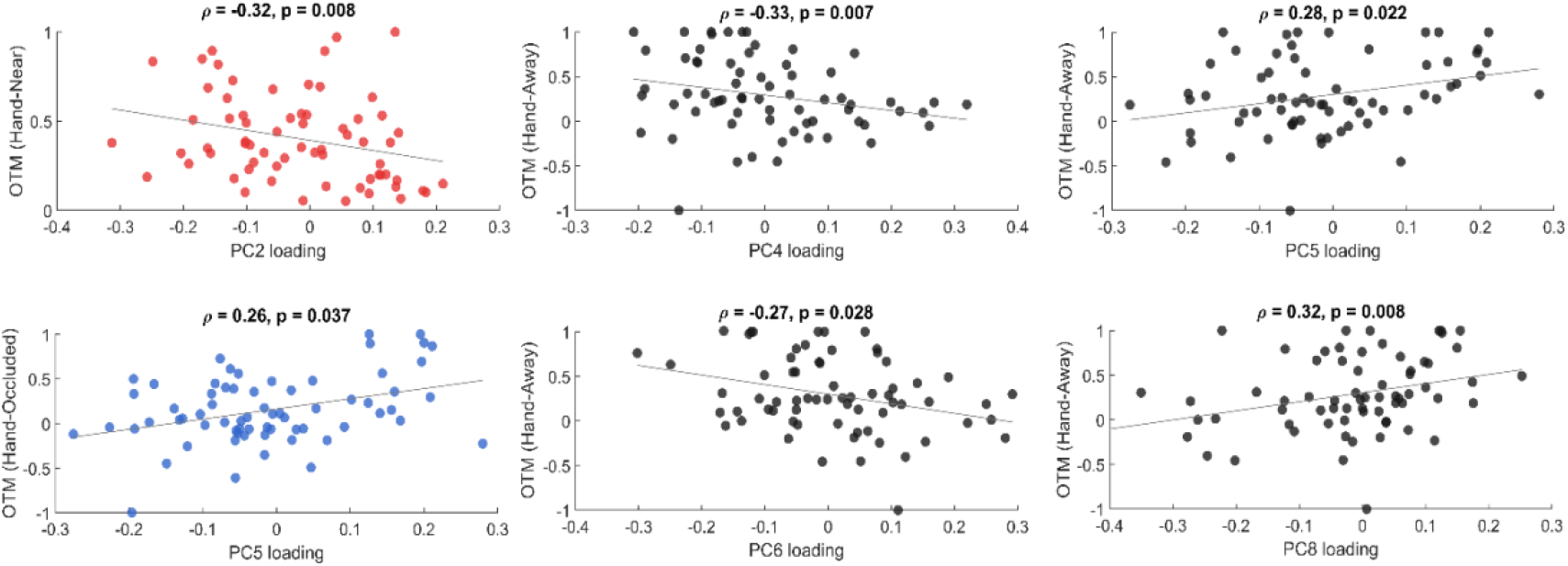
Correlations between orientation tuning modulation indices (OTM) and principal components (PCs).

